# Exercise-induced extracellular vesicles mediate apoptosis in human colon cancer cells in an exercise intensity-dependent manner

**DOI:** 10.1101/2024.09.24.614165

**Authors:** Berkay Ozerklig, Ibrahim Turkel, Merve Yılmaz, Refika Dilara Vaizoglu, Handan Sevim Akan, Z. Gunnur Dikmen, Ayesha Saleem, Sukran N. Kosar

## Abstract

Regular exercise is known to reduce incidence rates and improve the prognosis of all cancers, but the underlying mechanisms remain elusive. Ample evidence suggests that exercise exerts therapeutic effects through extracellular vesicles (EVs), essential for cellular communication. Here, we hypothesized that exercise-induced EVs from serum of healthy young male participants will exert anti-tumorigenic effects on human colon cancer HT-29 cells, in an exercise intensity-dependent manner. 10 healthy young active males (25.4±6.2yrs, with maximal oxygen consumption (VO2max) = 45±3.7 ml.min-1.kg-1 participated in a randomized crossover trial. Participants underwent two different workload-matched, acute bouts of exercise: (1) moderate-intensity continuous exercise (MICE) at 50-55% V02max, and (2) high-intensity interval exercise (HIIE) at 90% V02max on a cycle ergometer. A control session of rest (PRE) was included. EVs were isolated from serum samples collected during PRE and immediately after each exercise session. EVs were co-incubated with colon cancer HT-29 cells (100 µg EVs/ml, 48-72h), and effect on cell viability, migration, and apoptosis measured. EV treatment reduced cell viability in all groups (PRE, MICE, HIIE) by 35%, 43% and 47% respectively, vs. PBS. EVs from HIIE group showed a significantly greater reduction in cell viability vs. PRE, therefore further analysis used these groups only. PRE-EVs reduced migration by 27%, and HIIE-EVs by 39%. EV from HIIE group increased expression of pro-apoptotic markers: Bax/Bcl-2 ratio by 56% and Caspase-3 by 30% vs. PBS, with no change observed in the PRE group. Further, 16% of cells in PRE, and 28% of cells in HIIE were TUNEL-positive, indicating DNA fragmentation, a hallmark feature of apoptosis. Our data show that exercise-induced EVs reduced cell viability, in an exercise intensity-dependent manner. HIIE-derived EVs exerted the most anti-tumorigenic effects: decreased cell viability, reduced cell migration, increase in pro-apoptotic protein expression, and elevated DNA fragmentation. It is likely these changes were mediated by altered EV Cargo induced by exercise, as the amount of EVs was the same in each treatment group. To our knowledge, this is the first human study that illustrates the therapeutic potential of exercise-induced EVs in cancer treatment.

## Introduction

Colorectal cancer (CRC) is the third most common cancer worldwide and the second leading cause of cancer mortality, yet there is no cure for colorectal cancer to date [1-3]. Evidence suggests that regular exercise can be protective against CRC in a dose-dependent manner [4, 5], and is associated with decreased cancer-related mortality and recurrence [6]. Preclinical *in vivo* and *in vitro* studies demonstrate that systemic responses to exercise can inhibit cancer cell proliferation and metastasis [7, 8]. Exercise may exert these positive adaptations through the modulation of reactive oxygen species, inflammation, hormones, immune cell response, and growth factors, which ultimately attenuate tumor formation through alterations in the tumor microenvironment and circulating factors [9, 10]. However, the exact biological mechanisms underlying these anti-cancer protective effects of exercise have yet to be fully elucidated.

The beneficial systemic adaptations to exercise can be in part due to factors released as a physiological response to exercise. Growing evidence suggests that these exercise-induced molecules, named “exerkines”, such as cytokines, proteins, peptides, metabolites, and microRNAs (miRNAs) are secreted from metabolically active tissues, and exert pro-metabolic effects on multiple organs and biological processes in an autocrine, paracrine, and endocrine manner [11-13]. It follows that the transportation of these molecules to recipient tissues is essential for cellular communication and organ-organ crosstalk. Therefore, the mediators of such transport are pivotal. Exerkines can be released into circulation by themselves, or can be packaged within extracellular vesicles (EVs) [14, 15].

EVs are released from all cells, are essential to cellular communication and contain biomolecular cargo that can affect recipient cell function. EVs are divided into different groups according to their sizes and classically labelled as exosomes (30-150 nm), microvesicles (100-1000 nm), and apoptotic bodies (500-5000 nm) [16]. However, the Minimal Information for Studies of EVs (MISEV) 2023 guidelines recommend classifying EVs based on size (small EVs as <200 nm, and large EVs as >200 nm), composition, and cell of origin [17]. Acute exercise increases the circulating EV concentration and modifies EV cargo, in both rats and humans [18, 19]. EVs isolated from healthy endothelial and stem cells have therapeutic effects on different disease models *in vivo* and *in vitro* such as, ischemia-reperfusion injury in heart muscle [20], Alzheimer’s disease [21], chronic kidney disease [22], and CRC [23]. However, studies investigating the beneficial effects of exercise-induced EVs are sparse. A recent report showed that exercise-induced EVs attenuated tumor progression and metastasis in a sedentary syngeneic prostate tumor-bearing rat [24]. Similarly, another recent study demonstrated that EVs isolated from murine skeletal muscle myotubes after chronic contractile activity, an in vitro model of exercise, perpetuated anti-tumorigenic effects in Lewis Lung Carcinoma cells [25], and had an altered EV proteomic cargo profile [26].

Here, we investigated the effects of exercise-derived EVs from healthy young male participants in HT-29, a human colorectal adenocarcinoma cell line. We hypothesized that EVs isolated from participants immediately after acute exercise would inhibit cell growth by inducing apoptosis in HT-29 cells, in an exercise intensity-dependent manner. To evaluate this hypothesis, we used a randomized crossover experiment design with two different workload matched exercise models: moderate-intensity continuous exercise (MICE), and high-intensity interval exercise (HIIE). Blood samples were collected at PRE, post-MICE, and post-HIIE time points, and serum extracted for EV isolation and analysis. EVs isolated from 250 µl serum samples were co-cultured with HT-29 cells (100 µg/µl, 48-72h), and the effect on cell growth, viability, and apoptosis in recipient cells was measured.

## Methods

### Participants

Ten healthy recreationally active males (age: 25.4± 6.2 years) who engaged in physical activity 2-3 times per week were recruited. Participants were excluded if they had any known acute or chronic disease, smoking, or were taking any drug or supplements known to affect metabolism. Participants were given both oral and written information about the study procedures before they provided their written informed consent. The study was approved by the Hacettepe University Institutional Non-Interventional Ethical Committee with decision number 2022/20-58 and conducted in accordance with the Helsinki Declaration.

### Experimental design

Using a randomized crossover design, participants were randomly assigned to the MICE and HIIE groups for the initial intervention. Participants visited the laboratory four times, each visit separated at least 48-72h. On the first visit, body composition was measured, and a familiarization session was performed to accustom subjects to the equipment, maximal oxygen consumption (VO2max) test, and exercise interventions (MICE and HIIE). On the second visit, participants performed a progressive incremental exercise test on a bicycle ergometer to determine their respective VO2max. On the third and fourth visits, each participant performed a single session of either MICE or HIIE. Venous blood samples were collected at three time points: (1) immediately before the first exercise session (PRE), (2) immediately post-MICE session, and (3) immediately after HIIE session, and used for serum and subsequently EV isolation as described below. To minimize the influence of circadian rhythms, all exercise intervention protocols were performed at the same time of day (13:00 - 17:00h). Participants were instructed to maintain their usual exercise and dietary habits, but prior to the acute MICE and HIIE exercise sessions, participants were instructed to fast overnight, abstain from coffee and alcohol, and avoid strenuous exercise for 24h. Upon arrival at the laboratory, participants were asked to consume a 600 kcal breakfast and performed a single MICE or HIIE session 2h after breakfast.

### Body composition

The body composition of each participant was measured using dual-energy x-ray absorptiometry (DXA, Lunar Prodigy Pro Narrow Fan Beam (4.5°), GE Health Care, Madison Wisconsin, USA) following standardized procedures [27]. Body composition was measured after overnight fasting and body fat mass (kg), body fat (%), lean body mass (kg), and fat-free mass (kg) were determined.

### Determination of maximal oxygen uptake

All participants underwent an incremental exercise protocol on a bicycle ergometer (COSMED E200, Italy) using breath-by-breath technology (Quark CPET, Cosmed Cardio-Pulmonary Exercise Testing, Italy) to determine individual VO2max. Briefly, the exercise protocol started with a load of 60 W, which was increased by 30 W every 2 min. After the completion of three phases at 60, 90, and 120 W, the load was further increased by 30 W per min. Throughout the protocol, participants were instructed to maintain a stable, self-selected cadence of approximately 70 rpm for the duration of the test [28]. The test was terminated when at least two of the following criteria were met: (1) oxygen levels plateaued even with increasing workload, (2) the respiratory exchange ratio exceeded 1.1, (3) the heart rate was within 10 beats per min of the predicted maximum (220 minus age beats per min), and (4) unable to maintain 70 rpm for 10 seconds despite verbal encouragement [29].

### High-intensity interval exercise (HIIE) protocol

Although the literature lacks a clear approach on standardizing the parameters for the two different exercise protocols, previous work suggests equalizing the total workload (total exercise volume) and energy expended (isocaloric) [24]. After a 3 min warm-up at 60 W, participants performed 10 × 1min intervals at 90% of VO2max, separated by 75 seconds of active recovery at 60 W and a final 3 min cool down period at 60 W [30, 31].

### Moderate intensity continuous exercise (MICE) protocol

Similar to the HIIE group, participants performed a single session of MICE matched to the HIIE workload under supervision on a bicycle ergometer. Based on power output during HIIE session, the MICE group completed the ∼21 min of exercise at 55% of VO2max consisting of a 3 min warm-up and 3 min cool down at 60 W.

### Blood sampling and EV Isolation

Venous blood samples were collected from antecubital vein before and after the completion of each acute exercise session as described above in serum separating tubes (BD Vacutainer). Approximately 5 mL of blood was collected, allowed to clot for 40 min, and then centrifuged at 2000 x*g* for 10 min at 4°C, with resulting supernatant collected as serum. Serum samples were aliquoted and stored at -80°C until EV isolation. EVs were isolated from serum samples using ExoQuick Serum Exosome Precipitation Solution (System Biosciences, Palo Alto, CA, USA) according to manufacturers’ instructions and as before [32]. EV preparations were aliquoted and conserved at − 80°C for later use.

### NTA measurement with Nanosight NS300

EVs were characterized biophysically using the NanoSight NS300 Instrument (405 nm laser diode) according to the manufacturer’s protocols (Malvern Instruments Inc.), and in line with MISEV guidelines [17, 33]. The instrument was calibrated prior to each experimental run for nanoparticle size and quantity using standardized nanoparticle dilutions provided by the manufacturer. Suitable dilution of isolated EVs was defined before each measurement. EVs diluted by 1:1000 in sterile PBS and analyzed by NanoSight NS300. Each experimental run was performed in quintuplicate (each capture 60 seconds) and PBS was used to assess background. For each sample, the nanoparticle size distribution curve, refractive index, and the relative nanoparticle concentration of a particular size was recorded, with the cumulative percentage of nanoparticles.

### Transmission electron microscopy (TEM)

10 μl of 1:1000 diluted EVs in PBS were fixed with an equal volume of 4% paraformaldehyde and carefully applied onto a copper grid for 20 min at room temperature. Grids were washed with 100 μl PBS and fixed with 1% glutaraldehyde for 5 min. To remove the excess fixation buffer, the grids were washed 8 times with 100 μl distilled water, after which they were negatively stained with 1% uranyl acetate for 2 min and left to air dry at room temperature. TEM grids were visualized using a JEM-2100F Field Emission Electron Microscope operated at 80 kV using the fee-for-service at METU Central Lab (R&D Department TEM Laboratory, Middle East Technical University).

### EV protein extraction and quantitation

EV preparations were lysed using RIPA solution with protease inhibitor tablet (Roche) and EV concentration was determined using Pierce™ BCA protein assay kit (ThermoFisher Scientific) following manufacturer’s instructions and as before [34]. Briefly, the working reagent was prepared by mixing 24 parts of Reagent A, and 1 part of Reagent B. Standards were prepared by serial dilution of 2 mg/ml bovine serum albumin (BSA) ampule into clean vials using ddH_2_0. 25 μl of each standard or sample was added to a 96-well microplate in duplicates, followed by the addition of 200 μl of working reagent to each well. Samples were incubated at 37°C for 30 min and absorbance was measured at 562 nm using a microplate spectrophotometer (SpectraMax i3x, Molecular Devices, USA).

### Cell culture and EV treatment

HT-29 human colon cancer cells were a gift from Professor Z. Gunnur Dikmen (Hacettepe University) and originally purchased from American Type Culture Collection (ATCC, HTB-38). Cells were cultured in growth media made with fresh Dulbecco’s Modification of Eagle’s Medium (DMEM; Sigma-Aldrich) supplemented with 10% fetal bovine serum (FBS; Gibco/ThermoFisher Scientific) and 1% penicillin/streptomycin (P/S). Cells were grown at 37°C in 5% CO2 incubator for 24h. Cells were seeded in 96-well microplates for the MTT assay, and 24-well or 6-well plates for all other experimental outcomes. Cells were incubated at 37°C in a humidified atmosphere containing 5% CO2 for 24h before proceeding with EV treatment, as described below for each outcome variable.

### Assessment of cell viability

The effect of EVs on cell proliferation and viability were assessed by the colorimetric 3-(4,5-dimethylthiazol-2-yl)-2,5-diphenyltetrazolium bromide (MTT) assay as described previously [35]. MTT was purchased from Millipore Sigma (Milan, Italy). For this assay, 9,000 cells/well were seeded in a 96-well plate and co-cultured with 100 µg/µl EVs for 48h. MTT solution (1 mg/mL) was added to the cell medium, and cells incubated for 2h at 37°C in the dark. MTT is metabolized to formazan salt in viable cells [36]. Thus, cell viability was determined by analyzing the absorbance of the formazan read at 490 nm wavelength using a microplate spectrophotometer (SpectraMax i3x, Molecular Devices, USA).

### Cell scratch assay

Cell migration was assessed using a scratch wound healing assay [37]. HT-29 cells (2×10^5^ cells/well) were seeded in 24-well plates to grow in a monolayer for 24h. After treating the cells with EVs (100 µg/ml) a sterile 20–200 μL pipette tip was held vertically to scratch a cross in each well. The scratch closure was monitored and imaged in 0, 24h and 48h after scratch. The gap distance was quantitatively evaluated using ImageJ (National Institutes of Health). Cell migration was measured without inhibiting proliferation.

### HT-29 cell protein extraction, western blotting

To evaluate the anti-tumorigenic effects of exercise-EVs, we focused on the expression of intrinsic apoptosis pathway related proteins such as Bax, Bcl-2 and Caspase 3 [38], as well as an early marker of DNA double-strand breaks, γH2AX^Ser139^ [39]. Additionally, markers of EV sub-types were analyzed in line with international standardized guidelines for EV research [17]. Briefly, HT-29 cells were trypsinized and seeded at 6 well plates (6×10^5^ cells/well). Upon reaching to 80% confluence, cells were gently washed with PBS, and treated with 100 µg/ml EVs in fresh media for 72h. A longer time point was used to precisely evaluate the changes in protein expression and apoptotic induction. After 72h cells were lysed using ice cold RIPA buffer with protease/phosphatase inhibitor (Roche) for 30 min and centrifuged for 15,000 *xg* for 30 min. Supernatants were transferred in a new tube and protein concentration was determined using Pierce™ BCA protein assay kit (ThermoFisher Scientific) following the manufacturer’s instructions.

10-30 μg of total protein from HT-29 cell extracts were resolved on a 10-12% SDS-PAGE gel and subsequently transferred to PVDF membranes. Membranes were then blocked for 1h with 5% skim milk in 1xTris-buffered saline-Tween 20 solution (TBST) at room temperature, followed by incubation with primary antibodies in 3% skim milk overnight at 4°C. The following primary antibodies were used: rabbit monoclonal anti-CD63 (ab275018, Abcam, 1:1000) and TSG101 (ab275018, Abcam, 1:1000) for EV characterization. Rabbit polyclonal anti-Bax (A20227, ABclonal, 1:1000), anti-Bcl-2 (A0208, ABclonal, 1:1000), anti-Caspase-3 (A0214, ABclonal,1:1000), and anti-p-H2AX(Ser139) (#9718, Cell Signaling Technologies, 1:1000). Subsequently, membranes were washed three times for 5 min with TBST, followed by incubation with anti-rabbit (AS014, ABclonal) IgG HRP conjugated (1:1000-10000) in 3% skim milk for 1h at room temperature. After the membranes were washed, a chemiluminescence system (ChemiDoc™, Bio-Rad Laboratories) was used to detect labeled proteins. Quantification of the bands from the immunoblots was performed using computerized densitometry software ImageJ (National Institutes of Health, Bethesda, MD).

### TUNEL assay

TUNEL detection kit (Elabscience, E-CK-A322) was used for assessing DNA fragmentation, the hallmark measurement of apoptosis, after EV treatment. Briefly, 50,000 cells/well were seeded in 24-well plates and allowed to adhere for 6h, then co-incubated with 100 µg/ml EVs in fresh media for 72h. Treated HT-29 cells were fixed with 4% paraformaldehyde for 15 min and incubated with 1% Triton X-100 for 10 min at 37°C. After TUNEL staining, HT-29 cells were immersed in DAPI solution to stain cell nuclei, and washed three times with PBS. We used an Olympus (Olympus, Japan) IX70 inverted fluorescence microscope to acquire the images. The images were analyzed by ImageJ (National Institutes of Health, Bethesda, MD).

### Statistical analyses

All data were analyzed using paired Student’s t-test or one-way ANOVAs. Multiple comparisons in the one-way ANOVA were corrected using Tukey’s post hoc test. Individual data points are plotted, with mean ± standard error of mean (SEM) shown as applicable. All graphs were created using GraphPad-Prism software (version 10.1.2, GraphPad, San Diego, CA, USA). Significance was set at p<0.05, Exact p values for significant or close to statistically significant results is shown. An n=4-10 was conducted for all experiments.

## Results

### Participant demographics

The overall study design is shown in **Fig 1**. All participants recruited were male, 25.4± 6.2 years old, with average BMI of 23.5 kg/m^2^. Average body fat was 19.6% and average lean mass was 60.3kg (**Table 1**). The participants had an average VO2max of 45 ± 3.7 ml/kg/min (**Table 1**), which is considered an above average/good score for males of this age [40]. Combined with the 19.6 ± 4.9% body fat that falls under the healthy range for this age group [41], it indicates that the participants were physically fit, with good cardiorespiratory fitness levels.

**Table 1.** Participant characteristics showing mean ± SD.

| Characteristics | (n=10) |
| --- | --- |
| Age | 25.4 ± 6.2 |
| Body Mass (kg) | 73.4 ± 6.1 |
| Height (cm) | 176.9 ± 6.9 |
| Body Mass Index (kg/m<sup>2</sup>) | 23.5 ± 1.2 |
| Lean Mass (kg) | 60.3 ± 7.3 |
| Fat Mass (kg) | 14.2 ± 3.7 |
| Fat-free Mass (kg) | 60.3 ± 7.3 |
| Body Fat (%) | 19.6 ± 4.9 |
| VO2max (ml/kg/min) | 45 ± 3.7 |

Abbreviations: VO2max, maximum aerobic capacity

**Figure 1.**
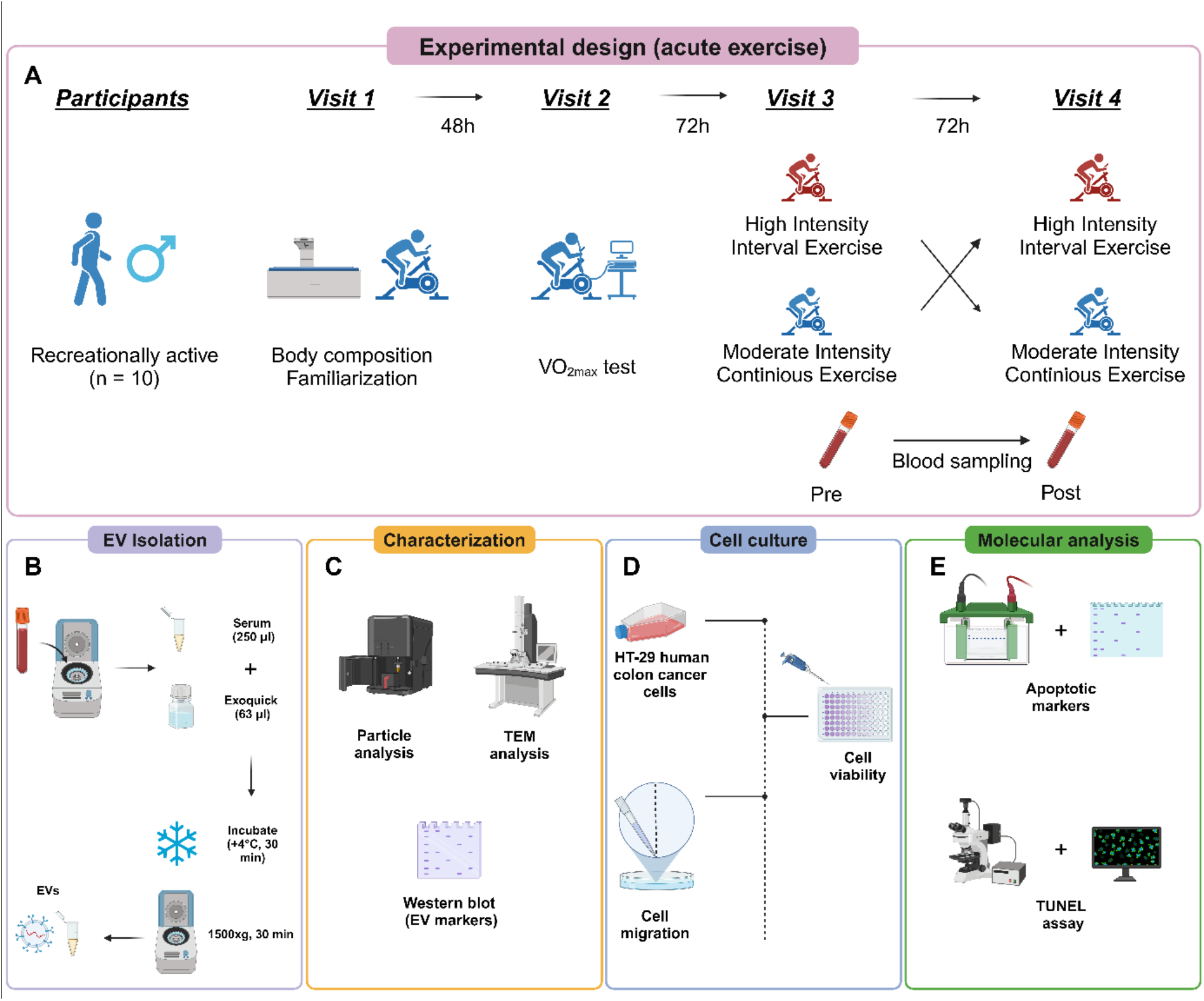
Experimental design of study. **(A)** Recreationally active *p*articipants were recruited and provided written informed consent. All participants attended the clinic four times. In the first visit, participants underwent a DXA scan to determine body composition and were familiarized with the equipment. In the second visit, a VO2max test was performed to calculate exercise intensities for HIIE or MICE intervention. Participants were then randomly allocated to HIIE or MICE groups, and completed each exercise intervention in the third and fourth visits as part of a randomized crossover design. Blood samples were taken immediately before (PRE) and after exercise tests (post). **(B)** EV isolation from serum samples of participants, **(C)** EV characterization with NTA, TEM, and western blotting was done in line with standardized guidelines for EV research. **(D)** Cell viability and cell migration experiments with HT-29 cells treated with EVs, as well as **(E)** western blot analysis for pro-apoptotic proteins and TUNEL assay for measuring DNA fragmentation in HT-29 cells treated with EVs were performed. Figure created using BioRender.com.

### Biophysical characterization of EV preparations and effect of exercise on small EV protein yield

Isolated EVs had an average size of 100.9 nm and an average concentration of 4.8E11 particles/ml (n=3, **Fig 2A**). TEM images demonstrated the isolation of intact double-membraned vesicles, roughly ∼100 nm in size (**Fig 2B**). Small EV markers TSG101 and CD63 were enriched in EV preparations (**Fig 2C**). EV depleted serum (EVds) was used as a negative control, and had significantly depleted expression of small EV protein markers. Altogether, EV characterization experiments illustrate that small EVs were successfully isolated. Finally, we noted that EV protein yield was increased in HIIE condition vs. PRE (p=0.0035, n=9, **Fig. 2D**).

**Figure 2.**
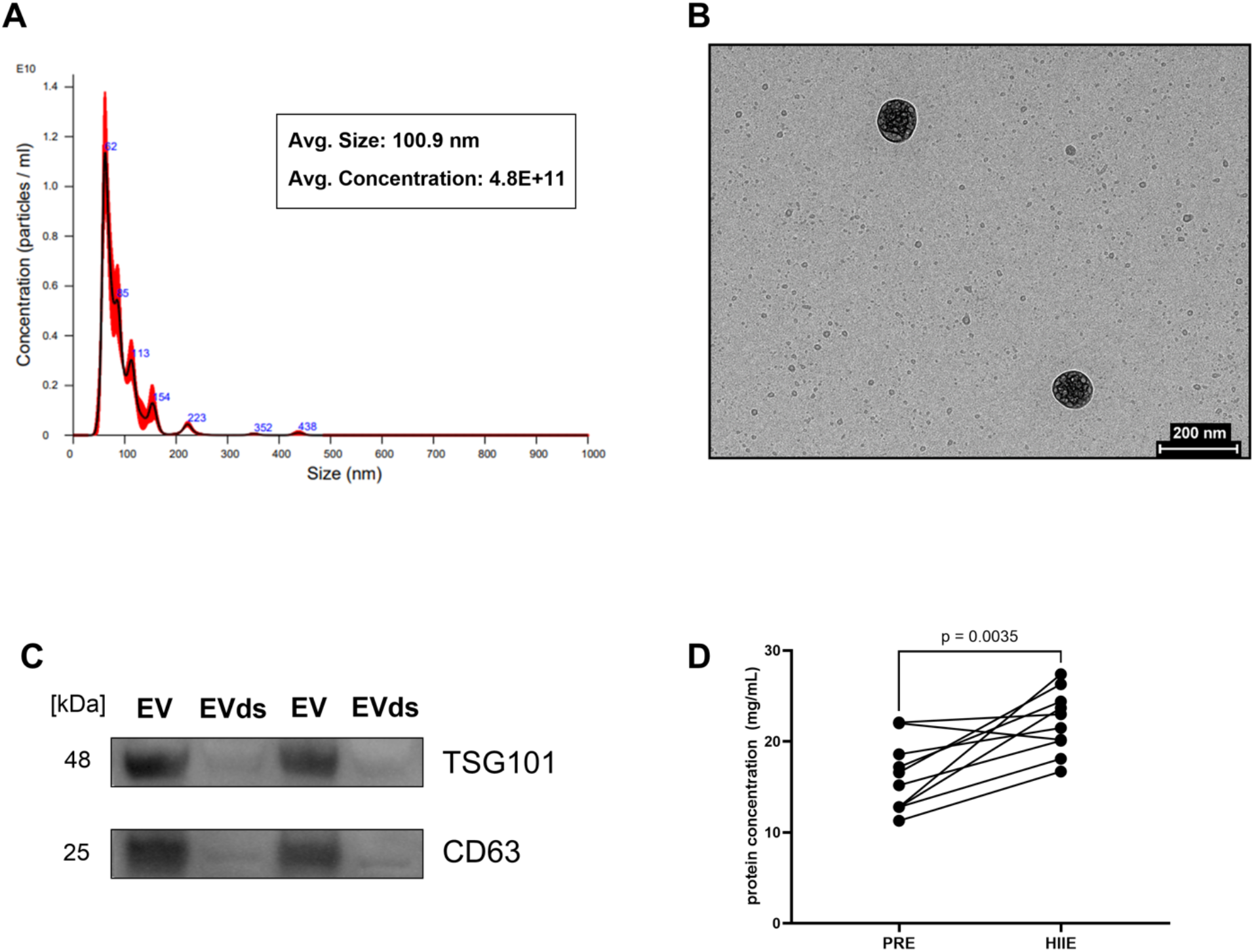
Characterization of EVs. EVs were isolated from 250µl serum and characterized using Nanoparticle Tracking Analysis (NTA), transmission electron microscopy (TEM), and Western blotting. **(A)** Representative NTA analysis graph showing size distribution and concentration of EVs isolated from human serum samples with average EV size and concentration data (n=3). We observed an enrichment of small EVs (<200 nm) in our EV preparations. **(B)** Representative TEM image, and **(C)** expression of small EV markers TSG101 and CD63 in isolated EVs and EV depleted (EVds) serum samples further confirmed the enrichment of small EVs in isolated EV fractions in the study. **(D)** Protein concentrations of EVs isolated from PRE and HIIE serum samples showed an increase in protein yield in HIIE group vs. PRE (p=0.0035, n=10). Data in 2D were analyzed using a paired Student’s t-test Significance was set at p<0.05.

### Exercise-induced EVs decrease cell viability and proliferation/migration in HT-29 cells

The effect of exercise-induced EVs on HT-29 cell viability was measured with an MTT assay. The results showed that cell viability decreased in all EV-treated groups compared to PBS: in PRE group by 35%, in MICE group by 43%, and in HIIE group by 47% (p<0.0001, n=5-10 per group, **Fig 3A**). Additionally, cells treated with EVs isolated from the HIIE group showed a 20% decrease in cell viability compared to cells treated with EVs from the PRE group (p=0.0480, n=5-10 per group, **Fig 3A**). As the HIIE group exhibited a significant reduction in cell viability compared to the PRE group, only this group was utilized as the exercise group for the remaining analyses. Next, we measured cell migration with a cell scratch assay, with the caveat that since cell growth was not controlled, the effects can be due to a combination of migration and cell proliferation. We did not observe any significant change after 24h of incubating the HT-29 cells with EVs (**Fig 3C**), however at the 48h time point cells treated with EVs from the PRE and HIIE groups showed decreased cell migration compared to PBS by 29%, (p=0.0416, n=4-6, **Fig 3D**) and 39%, respectively (p=0.0050, n=4-6, **Fig 3D**).

**Figure 3.**
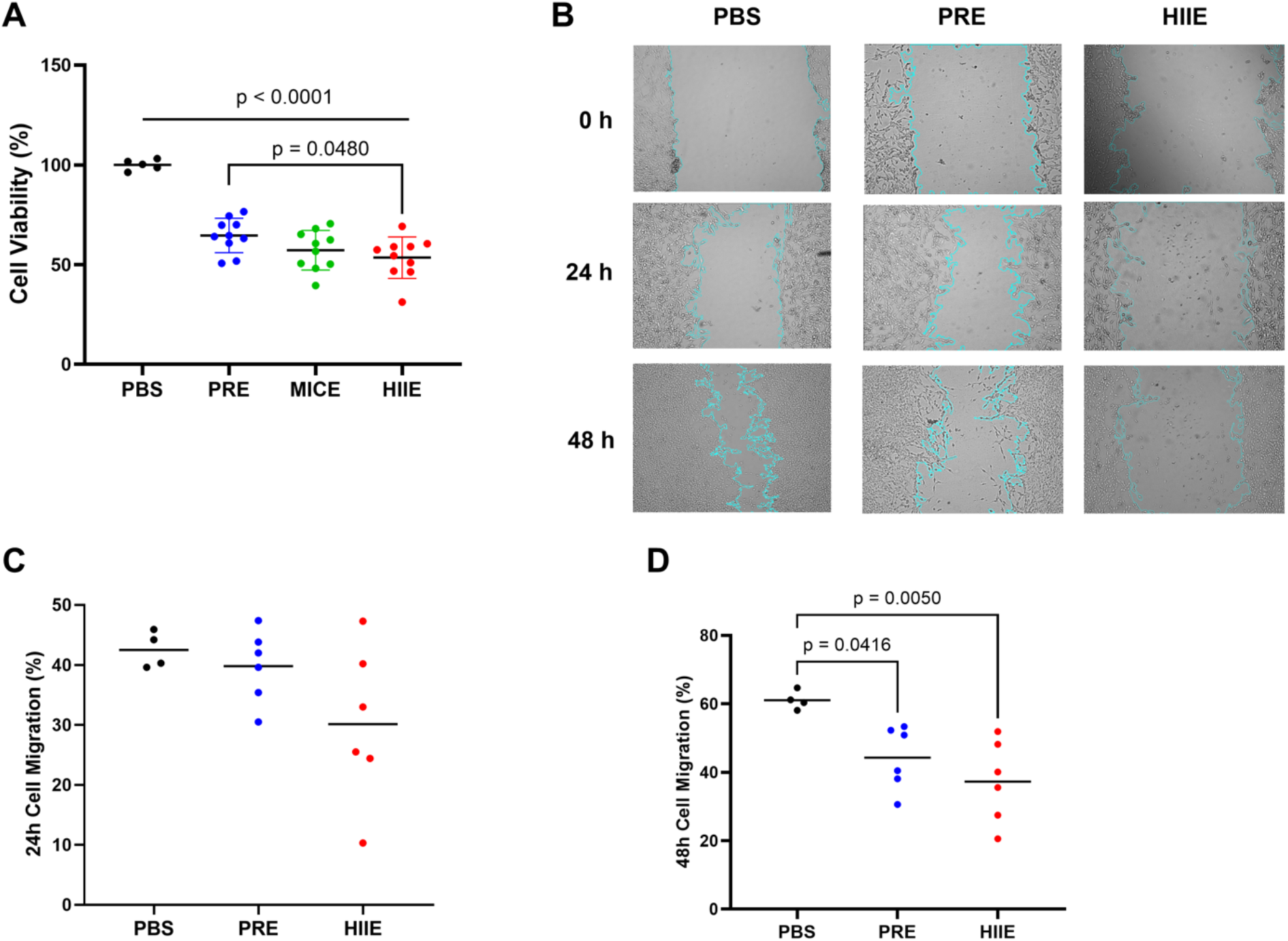
Cell viability and scratch assay. HT-29 colon cancer cells were treated with EVs (100 µg/ml) isolated from the different groups to elucidate if EVs could modulate cancer cell viability and migration. **(A)** Cell viability decreased in all EV-treated groups compared to PBS as determined using the MTT assay, in PRE by 35%, in MICE by 43%, and in HIIE by 47% (p<0.001, n=5-10 per group). Cells treated with EVs isolated from the HIIE group showed a further 20% decrease in cell viability compared to cells treated with EVs from the PRE condition (p=0.0480, n=5-10). **(B)** HT-29 cells were seeded in 24 well plates and kept in a 37°C CO_2_ incubator for 24h before EV treatment. Representative images from a cell scratch assay at 0, 24, and 48h post-PBS or EV treatment, with quantification of data in panels **(C)** and **(D). (C)** After 24h of EV treatment, there was no difference between groups. **(D)** After 48h, cells treated with EVs from the PRE and HIIE groups showed decreased cell migration compared to PBS by 29%, (p=0.0416, n=4-6), and 39%, respectively (p=0.0050, n=4-6). Data were analyzed using a one-way ANOVA with Tukey’s post hoc measurements. Significance was set at p<0.05. The p-values for non-significant data are not shown.

### Exercise-induced EVs induce apoptosis in HT-29 cells

To gain a deeper insight into the observed decrements in cell viability and migration with EV treatment, we measured expression of proteins involved in the intrinsic apoptotic pathway. Representative immunoblots for Bax, Bcl-2, Caspase 3, yH2AX (Ser139), and GAPDH as loading control are shown (**Fig 4A**), as well as quantification of results (**Fig 4B-4F**). While there were no statistically significant alterations in Bax (**Fig 4B**), and Bcl-2 (**Fig 4C**) expression with EV treatment, EV treatment from HIIE group increased Bax/Bcl-2 ratio by 57% (p=0.0252, n=6-10, **Fig 4D**), and Caspase 3 expression by 33% (p=0.0241, n=6-10, **Fig 4E**). yH2AX-Ser139 expression remained unchanged (**Fig 4F**).

**Figure 4.**
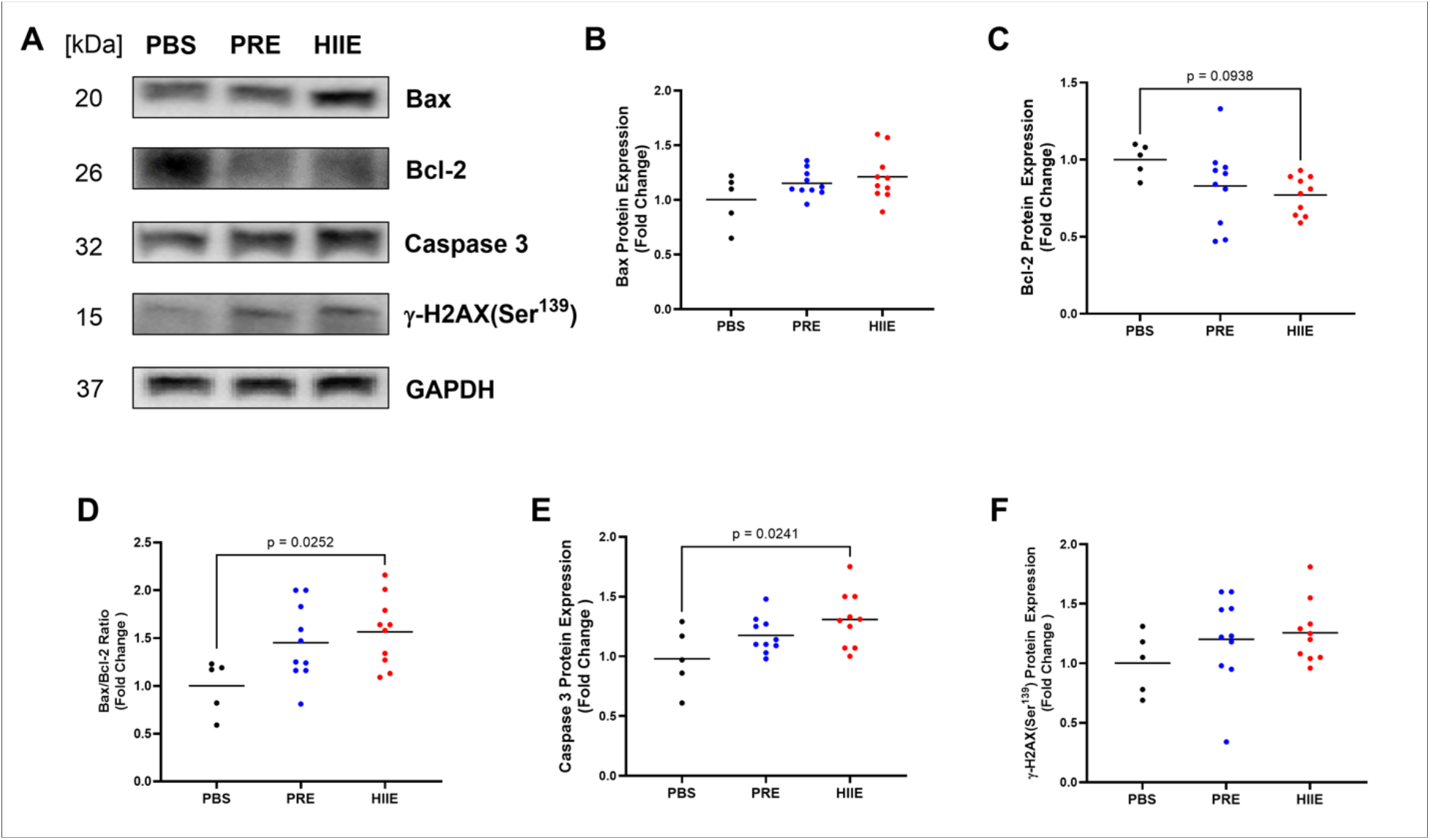
The expression levels of pro-apoptotic proteins in HT-29 cells after 72h of EV Treatment. Cells were treated with PRE and HIIE EVs for 72h, lysed, proteins extracted, and then resolved on a 10-12% SDS-PAGE for protein separation and Western blotting analysis. GAPDH was used as a loading control. **(A)** Representative immunoblots illustrating the expression of pro-apoptotic proteins in HT-29 cells treated with EVs from PRE, HIIE conditions, or PBS. There was no difference in protein levels of **(B)** Bax, **(C)** Bcl-2, and **(F)** γ-H2AX between the groups. **(D)** Bax to Bcl-2 expression ratio increased by 57% (p=0.0252, n=6-10), and **(E)** Caspase 3 expression increased by 33% in (p=0.0241, n=6-10) in cells treated with EVs from HIIE compared to PBS. Data were analyzed using a one-way ANOVA corrected with Tukey’s post hoc test. Significance was set at p<0.05. The p-values for non-significant data are not shown.

Finally, to ascertain if the HT-29 cells are undergoing apoptosis we measured DNA fragmentation, a hallmark feature of apoptosis, using a TUNEL assay. Representative images (**Fig 5A**) and quantification of data is shown (**Fig 5B**). EVs isolated from human serum samples increased TUNEL-positive nuclei by 1.6-fold in PRE group (p= 0.0062, n=6-8, **Fig 5B**) and by 2.8-fold in HIIE (p<0.0001, n=6-8, **Fig 5B**) compared to PBS. Altogether, exercise-induced EVs have a pro-apoptotic effect on HT-29 cells.

**Figure 5.**
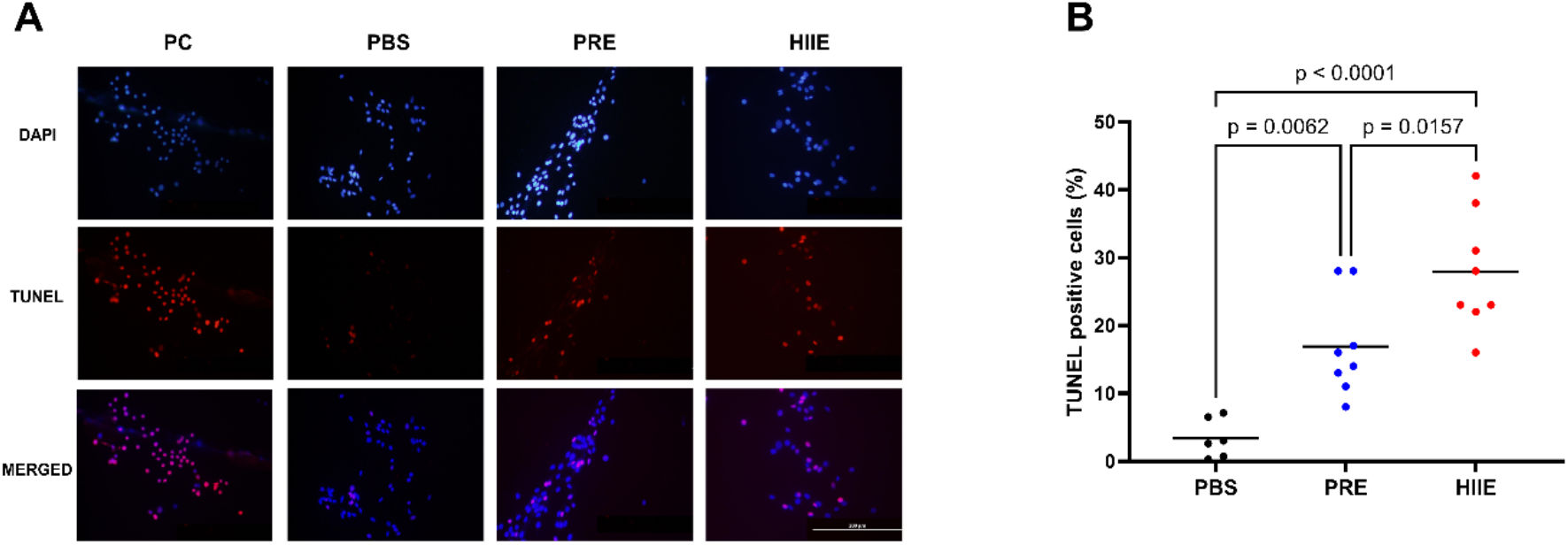
Evaluation of apoptosis in HT-29 cells treated with EVs using the TUNEL assay. HT-29 colon cancer cells were treated for 72h with EVs (100 µg/ml) isolated from PRE and HIIE conditions to elucidate if EVs could modulate apoptosis in cancer cells. DAPI was used to stain the cell nuclei. Scale bar = 200 µm. Apoptotic cells (TUNEL-positive, cells indicating DNA fragmentation, a hallmark feature of apoptosis) in four random fields were counted and used for apoptotic analysis. **(A)** Representative images of HT-29 cells incubated with EVs from PBS or PRE and HIIE groups for 72h. **(B)** EVs isolated from human serum samples induced apoptosis by 16% in PRE (p= 0.0062, n=6-8) and by 28% in HIIE (p<0.0001, n=6-8) compared to PBS. Data were analyzed using a one-way ANOVA corrected with Tukey’s post hoc test. Significance was set at p<0.05. The p-values for non-significant data are not shown.

## Discussion

Here we sought to investigate the effects of exercise-induced EVs on cell viability, migration/proliferation, and apoptosis in HT-29 human CRC cells. To our knowledge, this is the first investigation into the biological effects of human exercise-derived EVs on cancer cells. We found that exercise EVs, particularly those isolated from serum of health young males after high intensity exercise, reduced cell viability, mitigated migration/proliferation and induced intrinsic apoptosis culminating in increased DNA fragmentation, a hallmark measurement of apoptosis, in HT-29 cells.

We isolated EVs that were intact, enriched with small EVs as corroborated by EV size, TEM image, and protein marker expression data, in compliance with MISEV recommendations [17]. The measured EV concentration in our study was in line with estimated plasma EV concentrations as reported previously [42]. Interestingly we saw a 43% increase in serum EV protein yield in HIIE group compared to PRE group, in line with previous reports showing acute exercise increases EV concentration and/or EV protein yield [14]. For EV treatment, we selected an EV dose that seemed effective in previous work [4], and used the same dosage for each condition (PRE, MICE, HIIE). We recognize that given the increase in circulatory EVs with exercise in humans, the physiological dosage may be higher in vivo. EV treatment significantly reduced cell viability and migration/proliferation in all EV-treated groups, including the PRE EVs. A reduction in cell viability and migration/proliferation in the PRE group was unexpected. We posit that this is likely due to the above average cardiorespiratory fitness levels (average of ∼45 ml/kg/min VO2max, low body fat%) of our participants. We observed an additive decrease in cell viability, migration, and/or proliferation in the HIIE group vs. PRE, but not with MICE group, and subsequently, used PRE and HIIE groups for all further analyses. The exercise-intensity dependent effect is corroborated by previous reports where moderate-to-vigorous intensity acute exercise-conditioned serum exerts suppressive effects on different cancer cell lines including colorectal cancer cell lines [43]. It is postulated that this anti-tumorigenic effect is caused by an increase in exerkines, including SPARC, OSM, and IL-6 [43]. Cell migration is linked with cancer metastasis [44], thus inhibiting or attenuating cell migration is important for cancer treatment and this experiment is used frequently in cancer therapeutic research. Interestingly, we observed a significant reduction in cell migration in both PRE and HIIE groups at the 48h time-point, in parallel with our cell viability results. A recent study showed that exercise-conditioned serum derived from cancer patients inhibited the cell migration of pancreatic cancer cells [45]. Similarly, exercise-induced EVs isolated from rats attenuated the metastasis in sedentary tumor-bearing counterparts [46]. Since we did not inhibit cell proliferation while completing the scratch assay experiments, our results could be due to either migration or proliferation of cells. Future work where cell growth is inhibited by mitomycin C [25], can identify if the anti-migration effects are due to a reduction in cell movement or cell growth or both.

Next, we examined upstream protein mediators and end stage marker of apoptosis in CRC cells with EV treatment. Apoptosis is an essential programmed cell death pathway, and can be induced either intrinsically or extrinsically [47]. We observed a trend towards an increase in pro-apoptotic protein Bax content, concomitantly with a trend towards a decrease in anti-apoptotic Bcl-2 expression (p=0.09), with a significant increase in the overall Bax/Bcl2 ratio, a hallmark indicator of apoptosis, in the HIIE treatment group. Similarly, Caspase 3 expression increased significantly in HIIE group, which is well known initiator for DNA fragmentation in apoptosis. We measured γ-H2AX^(ser139)^ as a marker for double-strand DNA breaks, but did not see a significant increase with HIIE EV treatment. It is likely that a temporal increase in phosphorylation of γ-H2AX^(ser139)^, an early marker of DNA damage, was not captured in our experimental end time of 72h. Shorter time courses may reveal temporal changes in the induction of the upstream mediators of the intrinsic apoptotic pathway. Interestingly, similar to our cell viability/migration and western blot findings, we measured a significant increase in TUNEL positive nuclei, a hallmark marker for late-stage apoptosis, in both PRE and HIIE EV treated groups. Like with cell viability results, HIIE group had additive increase in the number of TUNEL-positive nuclei compared to PBS and PRE groups. Overall, it is interesting to see that exercise-induced EVs induce apoptosis in HT-29 cells.

Our study has a few limitations. First, we did not treat healthy non-cancer cells with the different EVs, so we cannot deduce if the HIIE EV treatment will reduce cell viability, migration/growth, and induce apoptosis in all cells or if the effect is specific to CRC cells. Two recent reports demonstrated that EVs released from C2C12 murine muscle myotubes following chronic contractile activity, an in vitro model exercise, induced a beneficial increase in mitochondrial biogenesis and elevated oxygen consumption rates [48] when incubated with C2C12 myoblasts, but induced cancer cell senescence, cell death and reduced viability when cultured with LLC cells [25]. This intriguing observation brings up a question as to whether exercise EVs might have different effects on different cell types, and the underlying mechanisms dictating recipient cell response to EV treatment. However, these questions require further investigation to elucidate the therapeutic effects of exercise EVs. Another limitation of this study is the level of training of the participants. A similar study in sedentary subjects would provide an opportunity to investigate the anti-tumorigenic effects of acute exercise-induced circulating EVs on cancer cells as a function of the fitness levels of the participating individuals. Additionally, the study was conducted only in young males, and the effects of sex/gender differences could not be examined. Conversely, given the average age of cancer patients, further investigation into the effects of exercise-induced EVs in the elderly is warranted.

In conclusion, our findings reveal for the first time that exercise-derived EVs from human participants reduce cell viability and induce apoptosis in HT-29 cells in an intensity-dependent manner. Further investigation of the EV cargo to identify biological factors, such as proteins, peptides, and miRNAs, will facilitate a deeper understanding of the therapeutic effects of exercise in prevention and potential treatment applications in cancer.

## Author Contributions

B.O., S.N.K., I.T. and A.S. contributed to the conception and design of the study. B.O., I.T., M.V., R.D.V., H.S.A performed experiments, analyzed the data, and created figures in the current study. B.O., A.S. and S.N.K helped write and revise the manuscript. B.O., I.T. A.S. and S.N.K. designed the project, and synthesize data, create figures, write, and edit the manuscript. All authors were involved in manuscript revisions.

## Acknowledgements

The authors would like to thank Suleyman Bulut for providing technical support and Beril Erdem Tuncdemir for assisting with immunoblotting of pro-apoptotic proteins.

